# Cryo-EM of human Arp2/3 complexes provides structural insights into actin nucleation modulation by ARPC5 isoforms

**DOI:** 10.1101/2020.05.01.071704

**Authors:** Ottilie von Loeffelholz, Andrew Purkiss, Luyan Cao, Svend Kjaer, Naoko Kogata, Guillaume Romet-Lemonne, Michael Way, Carolyn A. Moores

## Abstract

The Arp2/3 complex regulates many cellular processes by stimulating formation of branched actin filament networks. Because three of its seven subunits exist as two different isoforms, mammals produce a family of Arp2/3 complexes with different properties that may be suited to different physiological contexts. To shed light on how isoform diversification affects Arp2/3 function, we determined a 4.2 Å resolution cryo-EM structure of the most active human Arp2/3 complex containing ARPC1B and ARPC5L, and compared it with the structure of the least active ARPC1A-ARPC5-containing complex. The architecture of each isoform-specified Arp2/3 is the same. Strikingly, however, the N-terminal half of ARPC5L is partially disordered compared to ARPC5, suggesting that this region of ARPC5/ARPC5L is an important determinant of complex activity. Confirming this idea, the nucleation activity of Arp2/3 complexes containing hybrid ARPC5/ARPC5L subunits is higher when the ARPC5L N-terminus is present, thereby explaining activity differences between the different Arp2/3 complexes.

## Introduction

The Arp2/3 complex is evolutionarily conserved and built from two actin related proteins (Arp2 and Arp3) and five other protein subunits (ARPC1-5) (Goley and Welch, 2006; Molinie and Gautreau, 2018). It plays an essential role in many cellular processes, most notably cell migration (Goley and Welch, 2006; Krause and Gautreau, 2014), and also has newly identified roles in DNA repair (Caridi et al., 2018; Schrank et al., 2018). When activated by a nucleation promoting factor such as WAVE or WASP, the Arp2/3 complex is unique in its ability to assemble branched actin networks by stimulating new filament growth from the side of existing actin filaments (Campellone and Welch, 2010).

Until recently, the Arp2/3 complex has been considered as a single entity. However, in mammals, Arp3, ARPC1 and ARPC5 are present as two isoforms - Arp3, Arp3B, ARPC1A, ARPC1B, ARPC5 and ARPC5L - that share 91, 67 and 67% sequence identity respectively (Abella et al., 2016; Pizarro-Cerda et al., 2017). This raises the question as to whether different Arp2/3 complexes have evolved unique properties that are adapted to their particular cellular, developmental or physiological roles. Recent work has shown that the ARPC1 and ARPC5 isoforms differentially affect the actin nucleating properties of the Arp2/3 complex and the stability of the branched filament networks it generates (Abella et al., 2016). Furthermore, tissue-specific expression patterns of subunit isoforms, together with isoform-specific susceptibility to disease-causing point mutations, point to distinct physiological roles for particular Arp2/3 isoforms including cytotoxic T lymphocyte maintenance and activity (Brigida et al., 2018; Kahr et al., 2017; Kuijpers et al., 2017; Randzavola et al., 2019; Roman et al., 2017; Somech et al., 2017; Volpi et al, 2019).

Currently, the available high-resolution structures of mammalian Arp2/3 cannot address the role of isoform-specified diversity: using natively purified proteins, only a single isoform combination - ARPC1B-ARPC5 - of mammalian Arp2/3 is visualised in these structures (for example, the first structure described by (Robinson et al., 2001); this combination was shown to have intermediate nucleation activity (Abella et al., 2016). To begin to understand how subunit composition affects the properties of human Arp2/3 complexes, we used recombinant protein expression and cryo-electron microscopy (cryo-EM) to determine the structure of the most active human Arp2/3 complex, containing ARPC1B and ARPC5L subunits, referred to here as Arp2/3-C1B-C5L. We compared it with the structure of a complex containing ARPC1A and ARPC5 (Arp2/3-C1A-C5), which has the lowest activity (Abella et al., 2016). Our structures – the first sub-nanometre resolution reconstructions of any Arp2/3 complex determined by cryo-EM – show isoform-specific differences in the N-terminus of ARPC5/5L and suggest that these structural variations mediate different activities. Using protein engineering, we show that inclusion of the N-terminus of ARPC5L in hybrid Arp2/3 complexes enhances their actin nucleation activity, thereby showing how different Arp2/3 subunit isoforms contribute to differences in complex activation and function.

## Results & Discussion

### Overview of human Arp2/3 complex structure

We first determined the 4.2Å resolution structure and calculated a pseudo-atomic model of the most active human Arp2/3 complex Arp2/3-C1B-C5L (Figure 1; Table 1; Figure 1 – figure supplement 1) (Abella et al., 2016). The complex shows the characteristic triangle shape - ~150 Å ×130 Å and ~100 Å thick - seen in previous crystallographically determined structures of Arp2/3 complex purified from bovine brain and yeast (Nolen and Pollard, 2007, 2008; Nolen et al., 2009; Robinson et al., 2001). As observed in these previous structures, the intertwined ARPC2/4 subunits (Figure 1A,B, light/dark blue) form the platform for complex assembly, with Arp3/ARPC3 (Figure 1A,B, orange/magenta) and Arp2/ARPC1B/ARPC5L (Figure 1A,B, red/green/yellow)) constituting distinct protrusions.

**Table 1.** Model refinement statistics and geometries.

| **Model** | **Arp2/3-C1B-C5L** | **Arp2/3-C1A-C5** |
| --- | --- | --- |
| MolProbity score | 1.99 | 3.21 |
| Clash score | 11.1 | 46.89 |
| Bond mean RMSD (Å) | 0.007 | 0.018 |
| Angle mean RMSD (°) | 1.238 | 3.029 |
| Rotamer outliers (%) | 0.53 | 8.23 |
| **Ramachandran** |  |  |
| Outliers (%) | 0.30 | 0.85 |
| Allowed (%) | 6.2 | 4.49 |
| Favoured (%) | 93.51 | 94.65 |
| **Map/Model correlation coefficient** |  |  |
| CC (mask) | 0.78 | n/a (backbone only) |

**Figure 1.**
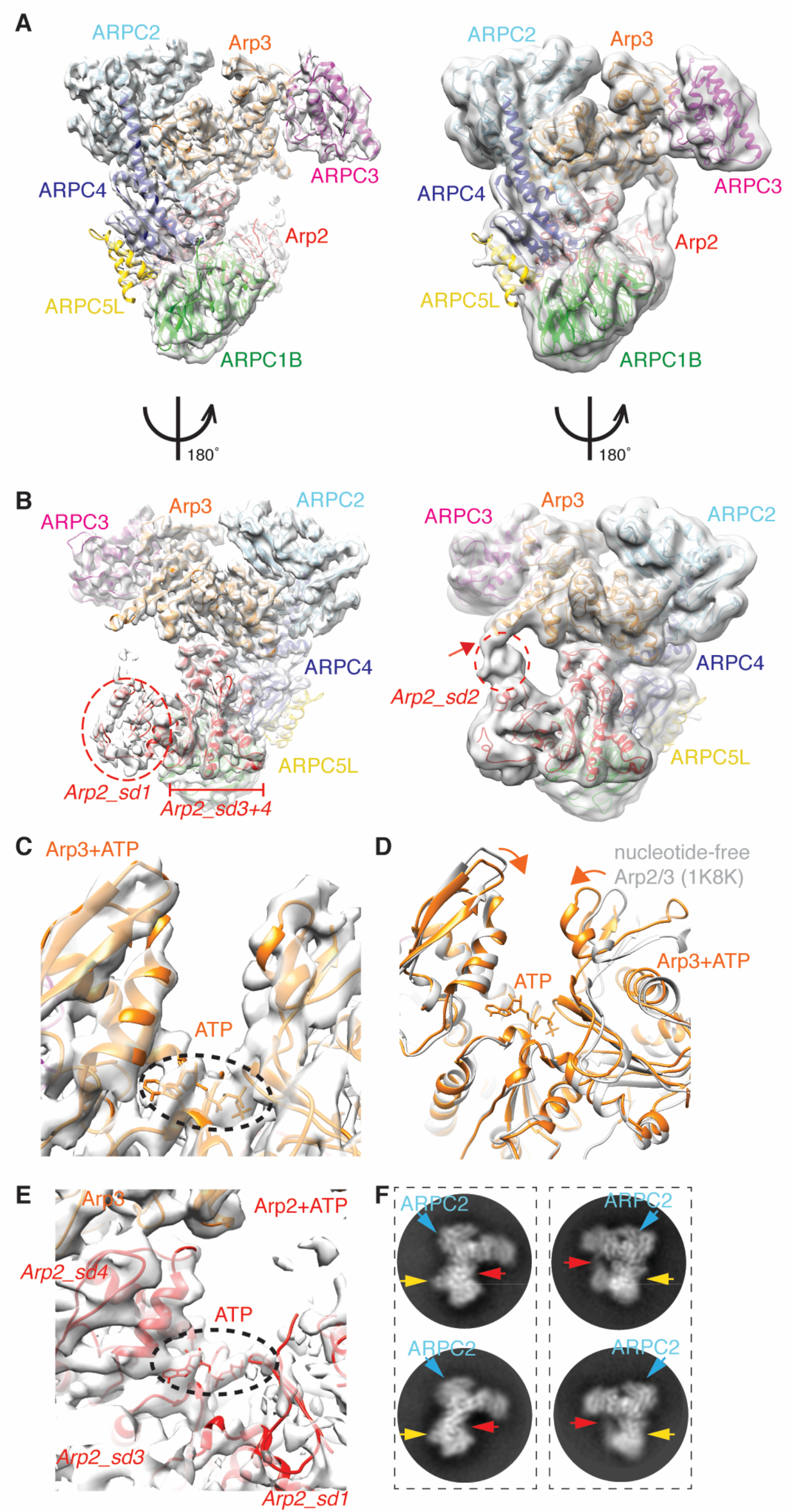
The cryo-EM structure of the human Arp2/3 ARPC1B-ARPC5L complex. (A) Left, Overview of the cryo-EM reconstruction of Arp2/3-C1B-C5L with the docked model in the density, viewed and coloured as originally presented by (Robinson et al., 2001): Arp2: red; Arp3: orange; ARPC1B: green; ARPC2: cyan; ARPC3: magenta; ARPC4: light blue; ARPC5L: yellow; Right, same view of the reconstruction with a ~8Å low-pass filter applied showing more flexible regions of the complex at lower resolution; (B) Left, 180º rotated view compared to (A) of the Arp2/3-C1B-C5L reconstruction and model; Right, same view of the reconstruction with a ~8Å low-pass filter applied showing more flexible regions of the complex at lower resolution, which includes flexible connectivity between Arp2 and Arp3 (red arrow) and parts of ARPC5L; (C) Cryo-EM reconstruction and model of nucleotide binding pocket of Arp3 with density corresponding to bound ATP indicated (dotted black oval); (D) Conformation of the nucleotide binding pocket of ATP-bound Arp3 in Arp2/3-C1B-C5L aligned (on subdomain 3) with a previously determined structure of nucleotide-free Arp2/3 (PDB 1K8K), showing closure of the pocket in the presence of bound nucleotide; (E) Cryo-EM reconstruction and model of nucleotide binding pocket of Arp2 with visible subdomain regions labelled and density corresponding to bound ATP indicated (dotted black oval); (F) 2D class averages of Arp2/3-C1B-C5L showing views corresponding to Figure 1A (left panels) and Figure 1B (right panels) illustrating the variable density corresponding to subdomain 2 of Arp2 (red arrows) and to ARPC5L (yellow arrows). ARPC2 is also labelled for reference (blue arrows).

Apart from a few residues at the N- and C-termini, the ARPC1B, C2, C3 and C4 subunits are completely visualised. Similarly, Arp3 is almost completely represented in the cryo-EM density. In contrast, while subdomains 3 and 4 of Arp2 are well ordered and at a similar resolution to the rest of the reconstruction, density corresponding to subdomain 1 is present but poorly defined and subdomain 2 is even less distinct (Figure 1B). In addition, although density corresponding to the 2 C-terminal α-helices of ARP5CL lies adjacent to ARPC4, there is a sharp fall-off in the strength of the cryo-EM density for the rest of the subunit further from the body of the complex (described in detail below). The regions of different flexibility within the complex also manifest in the variations in local resolution of the reconstruction (Figure 1 – figure supplement 1D).

### Conformation of ATP-bound Arp3 and Arp2

ATP is a key regulator of Arp2/3 complex activity (Ingerman et al., 2013; Le Clainche et al., 2001; Martin et al., 2006; Nolen and Pollard, 2007; Rodnick-Smith et al., 2016), and density corresponding to ATP is bound to Arp3 in the Arp2/3-C1B-C5L reconstruction (Figure 1C; Figure 1 – figure supplement 2A). Consistent with this, the Arp3 nucleotide binding pocket adopts a closed conformation as defined previously (Nolen and Pollard, 2007), such that the outer portions subdomains 2 and 4 of Arp3 are closer together than in structures with no bound nucleotide (e.g. 1K8K; Figure 1D). Furthermore, the conformation of human Arp3 in our reconstruction is similar to that of previously determined ATP-bound bovine Arp3 in this region (Figure 1 - figure supplement 2B).

In Arp2, density corresponding to bound nucleotide is present at the junction of subdomains 3 and 4 (Figure 1E; Figure 1 - figure supplement 2C). Density from Arp2 subdomain1 exhibits some ordering immediately adjacent to the nucleotide density, consistent with the role of this region in nucleotide binding, whereas only low-resolution density is present for most of subdomains 1 and 2, and is of insufficient quality for models to be calculated. The overall weak 3D density of this region of Arp2 is consistent with the conformational variability seen in 2D classes (Figure 1F), and with the flexibility that has been previously observed in Arp2/3 crystal structures (Nolen and Pollard, 2007, 2008; Nolen et al., 2009; Robinson et al., 2001). These data all support the idea that flexibility is intrinsic to this region of the Arp2/3 complex in the presence of ATP and that ATP binding does not induce large-scale conformational changes in the complex (Espinoza-Sanchez et al., 2018), in contrast to the large ATP-induced rearrangements previously proposed by (Rodnick-Smith et al., 2016). Earlier low-resolution EM characterizations of Arp2/3 probably also captured the dynamic properties of this region rather than activated states of the complex (Rodal et al., 2005; Sokolova et al., 2017). It is also possible that interaction with the grid surface in negative stain experiments may cause additional ordering of Arp2 subdomains 1 and 2 (Espinoza-Sanchez et al., 2018), in contrast to what we observe in our cryo-EM experiments.

The previously determined structure of inhibitor-bound Arp2/3 provides the most complete structural view of Arp2 (Luan and Nolen, 2013) in which subdomain 2 of Arp2 lies close to the loop connecting helix-α7 and -α8 in Arp3. Interestingly, although the Arp2 density in our uninhibited complex is relatively weak, the connectivity with Arp3 is not the same as in the inhibited structure, but rather occurs via helix-α9 (Figure 1 - figure supplement 2D). This supports the idea that Arp2/3 complex inhibition operates at least in part by restricting the movements of otherwise flexible regions of Arp2 and implies that flexibility in this region is important for subsequent steps in its activation. The connectivity between Arp2 and Arp3 in our reconstruction may represent a route of allosteric communication that is part of the activation process for the complex, a route that is blocked by inhibitor binding.

To investigate how general these observations are for cryo-EM-determined reconstructions of Arp2/3 complexes, we also calculated the 3D reconstruction of human Arp2/3 complex containing ARPC1A and ARPC5 subunits (Figure 1 - figure supplement 3A,B). This reconstruction has a slightly lower overall resolution (4.5 Å; Figure 1 - figure supplement 3C), its constituent particles also exhibit non-isotropic angular orientations (Figure 1 - figure supplement 3D) and the reconstruction shows non-isotropic variation in local resolution (Figure 1 - figure supplement 3E). However, it is clear from this structure that the overall conformation of the Arp2/3-C1A-C5 reconstruction is extremely similar to that of Arp2/3-C1B-C5L, apart from the density corresponding to ARPC5/C5L (see below). In particular, in each complex Arp2 and Arp3 are similarly organized with respect to each other, although subdomains 1 and 2 in Arp2 are even more flexible in the Arp2/3-C1A-C5 structure compared to Arp2/3-C1B-C5L (compare Figure 1 – figure supplement 3A,B,F with Figure 1A,B,F). This supports the idea that subdomains 1 and 2 of Arp2 are intrinsically flexible even in the presence of activating nucleotide, and that this flexibility is important for subsequent activation steps in the presence of additional ligands. The overall conformational similarity of these two isoform-specific complexes - which have been shown to have very different actin assembly promoting activities both *in vitro* and in cells (Abella et al., 2016) - implies that the isoform-specific differences observed are not mediated by fundamentally different architectures of the complex, but rather by alternative regulatory effects conferred by isoform-specific residues.

### Isoform specific subunit conformation and determinants of activity

To further investigate the differences between isoform-specific human Arp2/3 complexes, the location of partially- or non-conserved sequence variation between ARPC1A and ARPC1B were mapped onto the ARPC1B structure (Figure 2A, left, green spheres; Figure 2– figure supplement 1A). These isoform-specific-sites map primarily to the surface of the ARPC1 subunit rather than the β-propeller structural core (Figure 2A), consistent with the overall similar structures of each isoform. This suggests that Arp2/3 activity differences arising from ARPC1 isoforms are mediated either by the interactions with ARPC4 and ARPC5 within the complex itself, and/or with other binding partners, including the mother actin filament during branch formation.

**Figure 2.**
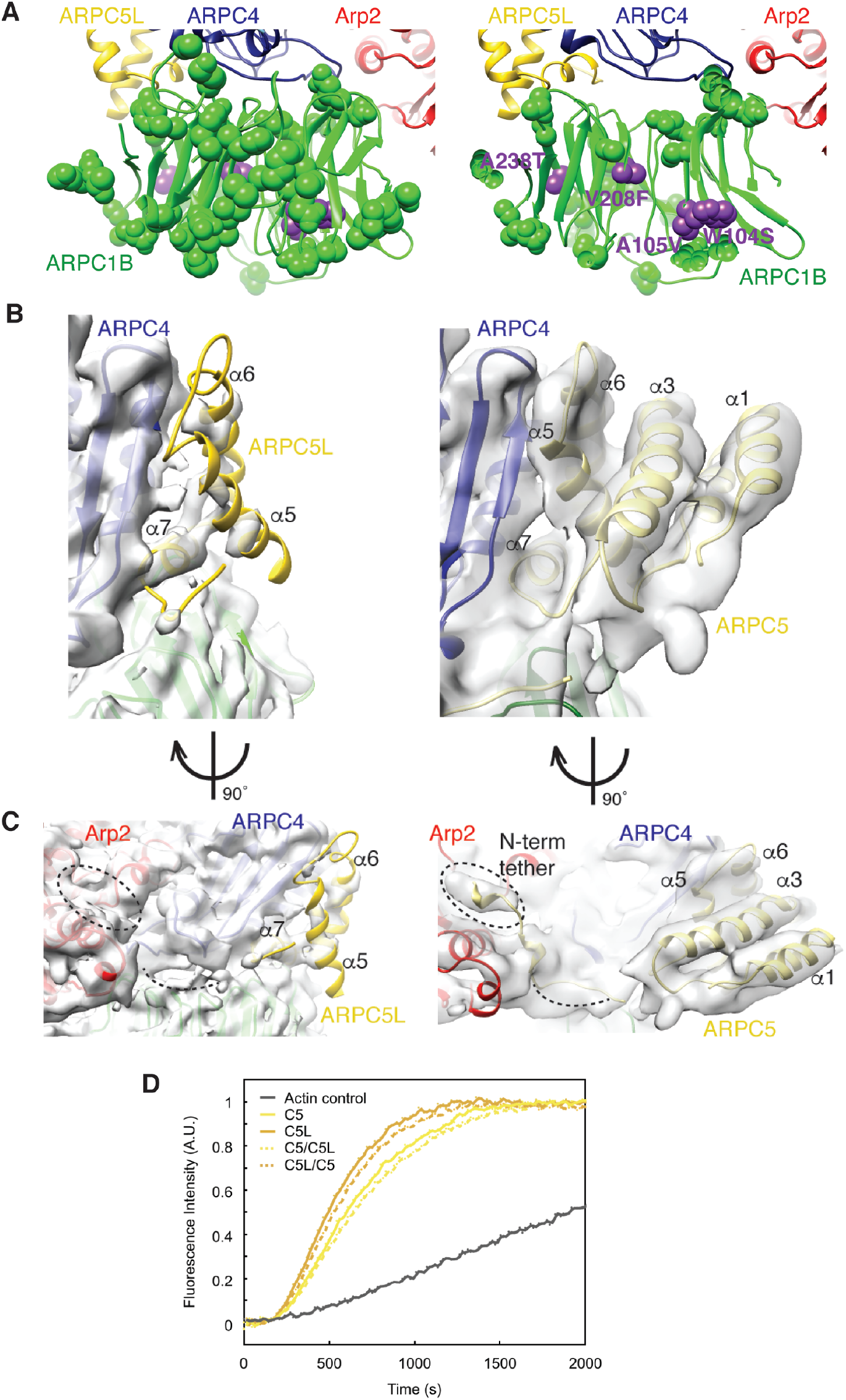
Isoform-mediated differences in human Arp2/3 complexes. (A) Left, location of non-conserved sequence variation between human ARPC1A and ARPC1B (green spheres) and location of disease-causing ARPC1B point mutations (purple spheres) mapped onto ARPC1B; right, cross section through ARPC1B; (B) Left, density corresponding to ARPC5L, showing the incomplete density for this subunit apart from helix-α7 adjacent to ARPC4; Right, density corresponding to ARPC5, showing the near complete density for this subunit (C) Left, 90º rotated view compared to B, left of ARPC5L showing the incomplete density for this subunit and lack of connectivity to Arp2; Right, 90º rotated view compared to B, right of ARPC5, showing the clear density for most of the subunit, including its N-terminal tether to Arp2. (D) *In vitro* polymerisation of 2μM pyrene-actin (5% labeled), either alone (black curve) or in the presence of 5 nM VCA and 1.25 nM of Arp2/3-C1A with the indicated ARPC5 isoforms or hybrids shows differences in actin assembly according to the ARPC5/5L N-terminal region present.

Loss-of-function mutations of ARPC1B give rise to multisystem inflammatory and immunodeficiency diseases (Brigida et al., 2018; Kahr et al., 2017; Kuijpers et al., 2017; Randzavola et al., 2019; Somech et al., 2017; Volpi et al, 2019), similar to Wiskott-Aldrich syndrome that is caused by mutations in the Arp2/3 activator WASP (Bosticardo et al., 2009). While some patients carry mutations causing premature protein termination and total loss of ARPC1B protein, four point mutations – W104S, A105V, V208F and A238T – have also been observed in patients (Brigida et al., 2018; Kahr et al., 2017; Volpi et al., 2019) (Figure 2A, right, purple spheres). W104, A105 and V208 are located in the core of the ARPC1B subunit β-propeller fold and this mutation likely destabilizes its tertiary fold, causing severely reduced protein levels in patients (Kahr et al., 2017). Conversely, A238T is more peripheral in the structure indicating that more than one disease mechanism may be in play. Although some of these patients show increased levels of ARPC1A, the activity of this isoform is insufficient to substitute for lack of functional ARPC1B, consistent with the idea that Arp2/3 complex subunit diversity has been tuned for particular physiological contexts.

The biggest difference between our two Arp2/3 cryo-EM reconstructions is the ARPC5 subunit (Figure 2B). In the Arp2/3-C1A-C5 structure, density corresponding to the helices of ARPC5 is clearly visible (Figure 2B, right), as is the N-terminus of the subunit that extends to contact the barbed end cleft of Arp2 (Figure 2C, right). In the Arp2/3-C1B-C5L reconstruction, however, density corresponding to ARPC5L is overall much weaker (Figure 2B, left). Density corresponding to the ARPC5L C-terminal helix-α7 is present and forms the major contact with the body of the Arp2/3 complex via ARPC4; at lower thresholds, density corresponding to helix-α5 and –α6 are also visible (Figure 1B, right). However, the N-terminal half of this subunit, including connectivity to Arp2, is not visible even at low thresholds (Figure 2C, left). In principle, this could be due to variable ARPC5L protein occupancy in the complex, but the well-defined density of helix-α7 at the interface with ARPC4 suggests this is not the case.

The C-terminal portion of ARPC5 - visible in both isoform reconstructions - corresponds to regions of greatest sequence similarity between the isoforms and between organisms, whereas the N-terminal region is more divergent (Figure 2 – figure supplement 1B). Our structures suggest the intriguing possibility that the higher activity of Arp2/3-C1B-C5L complexes (Abella et al., 2016) may be due to the intrinsically more flexible attachment of ARPC5L N-terminus to the complex. To test this idea, we engineered ARPC5/C5L protein hybrids, in which their N-terminal regions were swapped (Figure 2 – figure supplementary 1B; figure supplement 2A-C), and tested their actin nucleation activity in the context of the lowest activity Arp2/3-C1A complex. Consistent with the prediction arising from our structures, the hybrid complex containing the N-terminal region of ARPC5L was more active than that containing the equivalent region of ARPC5 (Figure 2D).

We therefore suggest that while the C-terminus of ARPC5 is closely associated with the body of Arp2/3, the role of its N-terminus is to contribute to regulation of complex activation: when contacting Arp2, the complex is held in an inactive conformation, while priming of the complex involves release of these contacts and conformational restraints between the subunits. With its looser, less structured N-terminal connectivity, human ARPC5L-containing complexes are intrinsically more able to undergo activator-induced conformational changes, and may also adopt alternative conformations in the activated complex (Dalhaimer and Pollard, 2010). All currently available X-ray crystal structures of mammalian Arp2/3 complexes contain the ARPC5 isoform and most of these structures fully visualize ARPC5; however, some also exhibit ARPC5 N-terminal conformational variability. Together with lower resolution negative stain EM analysis of *S. cerevisiae* Arp2/3 complexes (Martin et al., 2005) and NPF-bound *S. pombe* Arp2/3 complex (Espinoza-Sanchez et al., 2018), these observations support the wider functional relevance of the dynamics of the ARPC5 N-terminus in Arp2/3 activation revealed in our cryo-EM reconstructions.

The overall conformations of our ATP-bound Arp2/3 cryo-EM structures are similar to previously determined X-ray crystallography structures of natively purified bovine Arp2/3 complexes (Nolen et al., 2004; Nolen and Pollard, 2007). Earlier lower resolution negative stain EM studies of Arp2/3 complex have described a range of conformational changes in the Arp2/3 complex in response to ATP binding that likely reflect the intrinsic flexibility of portions of the unactivated complex, especially within Arp2 (Rodal et al., 2005; Sokolova et al., 2017). Thus, rather than inducing major conformational changes in the complex, available structures of Arp2/3 together with biophysical measurements point to the role of ATP binding in potentiating subsequent conformational changes induced by other activating factors, including nucleation promoting factors and binding to the mother filament (Goley et al., 2004; Kiselar et al., 2007; Luan et al., 2018; Martin et al., 2005). Our results set the stage for cryo-EM structure determination of Arp2/3 complexes bound by nucleation promoting factors, promising insights concerning steps of Arp2/3 activation - including isoform-dependent modulation of complex activation - in the future.

## Materials and Methods

### Cryo-EM grid preparation and data collection

Arp2/3-C1B-C5L and Arp2/3-C1A-C5 complexes were prepared using the multibac expression system consisting of three pFL vectors (pFL–ARPC2–Arp3; pFL–ARPC4– ARPC1A or C1B–STREP; and pFL–ARPC3–ARP2–ARPC5 or C5L) as described previously (Abella et al., 2016). They were diluted to 0.2-0.3 mg/ml in MOPS buffer (20 mM MOPS pH 7.0, 100 mM KCl, 2mM MgCl_2_, 5 mM EGTA, 1 mM EDTA, 0.5 mM DTT, 0.2 mM ATP) and 4 μl were applied to glow-discharged 1.2/1.3 holey gold grids (Quantifoil) before plunge freezing in liquid ethane using a Vitrobot (FEI/Thermo Fisher Scientific) operating at room temperature and 100% humidity. Data for the Arp2/3-C1B-C5L complexes were collected at the UK National Electron Bio-imaging Centre (eBIC) at the Diamond Synchrotron using EPU on a Titan Krios microscope (FEI//Thermo Fisher Scientific) operating at 300 kV, and recorded on a K2 direct electron detector equipped with a Quantum energy filter (Gatan), with a pixel size of 1.06 Å/px. Two movies per hole were collected using a total dose of 60 e^−^/Å^2^ each over 20 frames with 8 s exposure, and with a defocus range of −1.5 – −3.5 μm. Data for the Arp2/3-C1A-C5 complexes were collected manually on a Polara microscope (FEI/Thermo Fisher Scientific) operating at 300kV, and recorded on a K2 direct electron detector equipped with a Quantum energy filter (Gatan), with a pixel size of 1.09 Å/px. A total dose of 59 e^−^/Å^2^ per movie over 33 frames with 10 s exposure, and with a defocus range of −1.5 – −3.5 μm.

For movie processing, all frames were aligned with using MotionCor2 (Zheng et al., 2017) and CTF was estimated with ctffind4 (Rohou and Grigorieff, 2015). Each dataset was processed independently using the RELION beta-version 2.0 (Kimanius et al., 2016). For the Arp2/3-C1B-C5L dataset, 1,547,329 particles were picked automatically from 3039 micrographs in RELION v1.4 and v2.0 (Kimanius et al., 2016; Scheres, 2012), with a box-size of 240 x 240 pixels. After iterative cleaning of the particles using 2D classification and particle sorting, 384,330 particles remained. These were subjected to a first 3D refinement using a crystal structure of the bovine ARP2/3 complex (PDB: 3DXM (Nolen et al., 2009) converted to EM density in EMAN (Ludtke et al., 1999) and filtered to 20 Å as an initial reference. The resulting initial reconstruction was used as a reference for several further rounds of 2D and 3D classification in RELION. The final 3D-reconstruction containing 101,606 particles, was calculated from dose-weighted data and was automatically B-factor sharpened in RELION using with a B-factor of −151 (Scheres, 2012). The final overall resolution of the masked reconstruction was 4.2 Å (0.143 gold-standard-FSC). For the Arp2/3-C1A-C5 dataset, a similar procedure was followed: 697, 272 particles were picked from 1221 micrographs with a box-size of 256 x 256 pixels. After iterative cleaning of the particles using 2D classification and particle sorting, 405,152 particles remained. Because of the highly preferred orientation of particles in this dataset, particles with over-represented orientations were removed by 2D and 3D classification. The final 3D-reconstruction containing 130,973 particles was calculated from dose-weighted data and was automatically B-factor sharpened in RELION using with a B-factor of −179 (Scheres, 2012). The final overall resolution of the masked reconstruction was 4.5 Å.

### Model building

An initial model of Arp2/3-C1B-C5L was created using coordinates from PDB file 1K8K, and was rigid-body docked to the cryo-EM map using Chimera (Pettersen et al., 2004). Additional elements including the nucleotides from PDB file 4XF2 were added, before the bovine sequences from these structures were altered to human. Secondary structure-based rigid bodies were described with Ribfind (Pandurangan and Topf, 2012), before a combination of conjugate gradient energy minimisation and molecular dynamics fitting of the model into the map was undertaken with Flex-EM (Topf et al., 2008), part of the CCP-EM suite (Burnley et al., 2017). The final model was processed using real space refinement in Phenix (Adams et al., 2010), for which a refinement resolution cutoff of 5 Å was used. Molprobity (Chen et al., 2010) was utilised to validate the geometry of the resultant model, which was then improved by manual inspection and local refinement of poor areas in Coot (Emsley et al., 2010), with Ramachandran and secondary structure restraints utilised. Because the Arp2/3-C1A-C5 reconstruction showed substantial non-isotropic resolution, a separate model for this structure was not calculated. Instead, the refined Arp2/3-C1B-C5L model was rigidly docked into the Arp2/3-C1A-C5 density, the ARPC5 model was replaced with that from 1K8K and its docking was locally refined. A full validation report for the atomic model was generated in phenix v1.17.1-3660 (Table 1).

### Production of ARPC5/C5L hybrid complexes

ARPC5/C5L hybrid-containing complexes were designed by replacing residues 1-95 of C5L with 1-93 of C5, while for ARPC5L/C5 residues 96-153 of C5L were replaced with 94-151 of C5 (Figure 2 – figure supplement 2A). DNA corresponding to the hybrids were obtained from GeneArt Gene synthesis (Thermo Scientific), which were cloned into pFL vector to obtain pFL-ARPC5/C5L using BamHI/NotI sites and pFL-ARPC5L/C5 using XhoI/XmaI sites, respectively. pFL-ARPC5/C5L was digested using BstZ17I/AvrII and the resulting insert was ligated into pFL-ARPC3-ACTR2 that was linearized using BstZ17I/SpeI, to create pFL-ARPC3-ACTR2-ARPC5/C5L. pFL-ARPC5L/C5 was digested using BstZ17I/PmeI and the insert was ligated into pFL-ARPC3-ACTR2 that was linearized with BstZ17I, to generate pFL-ARPC3-ACTR2-ARPC5L/C5. The expression in Sf21 insect cells and protein purification of ARP2/3 complexes containing the hybrids was performed as previously described (Figure 2 – figure supplement 2B) (Abella et al., 2016). To validate expression of the hybrids, Near-infrared Western immunoblot using Odyssey CLx detection system (Li-COR) was performed on the purified complexes with the following antibodies: Arp3 (Sigma A5979, Mouse), ARPC5L (Abcam ab169763, Rabbit), ARPC1A (Sigma HPA004334, Rabbit), and ARPC5 (Santa Cruz sc-166760, Mouse) (Figure 2 – figure supplement 2C).

### Actin nucleation assays

Recombinant GST-tagged VCA domain of human N-WASP (391-505) was expressed in E.Coli Rosetta 2 (DE3) and purified by affinity chromatography over a Sepharose 4B GSH affinity column (GE healthcare) followed by gel filtration over a Hiload Superdex 200 column (GE healthcare). Skeletal muscle actin was purified from rabbit muscle acetone powder following the protocol described in (Wioland et al., 2017). Pyrenyl-actin was made by labeling actin with N(1-pyrene)-iodoacetamide (Thermo Fisher).

Actin assembly was detected by the change in pyrenyl-actin fluorescence using a Safas Xenius spectrofluorimeter (Safas) at room temperature. 1 μL of 0.2 μM Arp2/3, 1 μL of 0.8 μM VCA and 8 μL of 20xKME (4 mM EGTA, 20 mM MgCl2, 1 M KCl) were mixed with 110 μL of G-buffer (10 mM Tris HCl pH 7.0, 0.2 mM ATP, 0.1 mM CaCl_2_, 1 mM DTT). 40 μL of 8 μΜ Mg-ATP-G-actin (5% pyrene labeled) was added to this protein solution and mixed rapidly. The fluorescence signal was recorded immediately, and until the curves reached the steady-state plateau. The fluorescence intensity was normalised using I(t)=(I_obs_(t)−I_min_)/(I_max_−I_min_) where I_min_, the average of the ten lowest data points, refers to the signal intensity before actin started to polymerise and I_max_, the average of the ten highest data points, refers to the signal intensity at steady-state. The experiments were repeated four times, giving similar results.

## Acknowledgements

OvL and CAM were supported by the Biotechnology and Biological Sciences Research Council (BB/L00190X/1), and acknowledge Diamond for access and support of the Cryo-EM facilities at the UK National Electron Bio-imaging Centre (eBIC), proposal EM14498, funded by the Wellcome Trust, MRC and BBSRC. We thank Szymon Manka for help with model calculations, the Birkbeck EM group for helpful discussions and Corey Hecksel, Alistair Siebert and Dan Clare at eBIC for assistance with data collection. MW is supported by the Francis Crick Institute, which receives its core funding from Cancer Research UK (FC001209), the UK Medical Research Council (FC001209), and the Wellcome Trust (FC001209). GRL was supported by the Agence Nationale de la Recherche (grant MuScActin). This project has received funding from the European Research Council (ERC) under the European Union’s Horizon 2020 research and innovation programme (grant agreement No 810207).

## Data Deposition

The cryo-EM maps and the corresponding structural coordinates were deposited under the accession codes PDB: 6YW6 and EMDB:10959 for Arp2/3-C1B-C5L and PDB:6YW7 and EMDB:10960 for Arp2/3-C1A-C5.

---

**Figure 1 - figure supplement 1.**
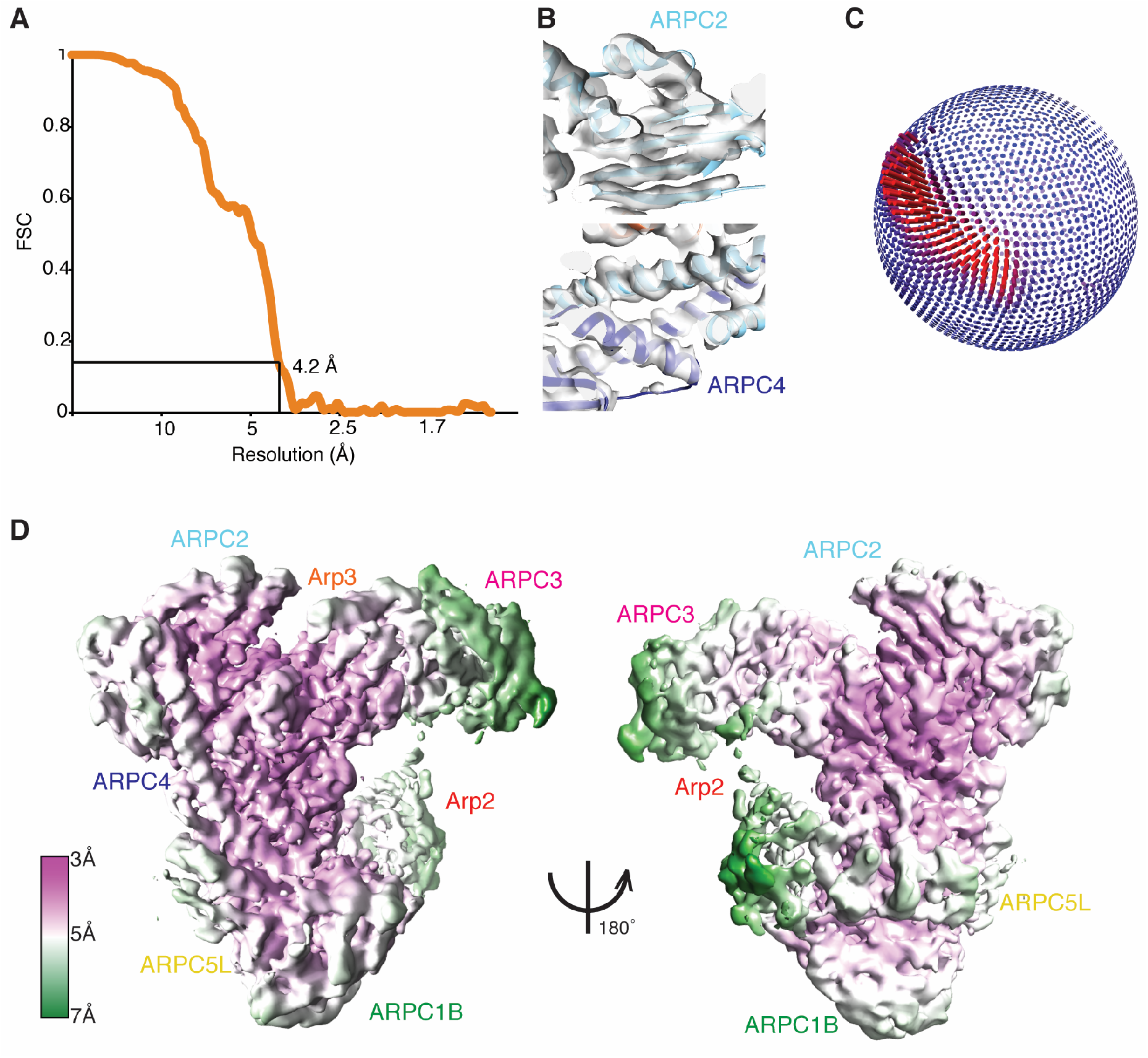
Evaluation of the resolution of the Arp2/3-C1B-C5L reconstruction. (A) FSC curve of the Arp2/3-C1B-C5L reconstruction, showing the 0.143 criteria resolution cut-off = 4.2 Å; (B) Example sections of the Arp2/3-C1B-C5L cryo-EM density showing a β-sheet region of ARPC2 (top) and α-helical regions of ARPC2 and ARPC4 (bottom) illustrating the quality of the best regions of the reconstruction; (C) Orientation distribution of particles used for the final 3D reconstruction; the most common view corresponds to that depicted in Figure 1A; (D) Local resolution depiction of the Arp2/3-C1B-C5L reconstruction calculated using Relion, showing views equivalent to Figure 1A (left) and Figure 1B (right).

**Figure 1 - figure supplement 2.**
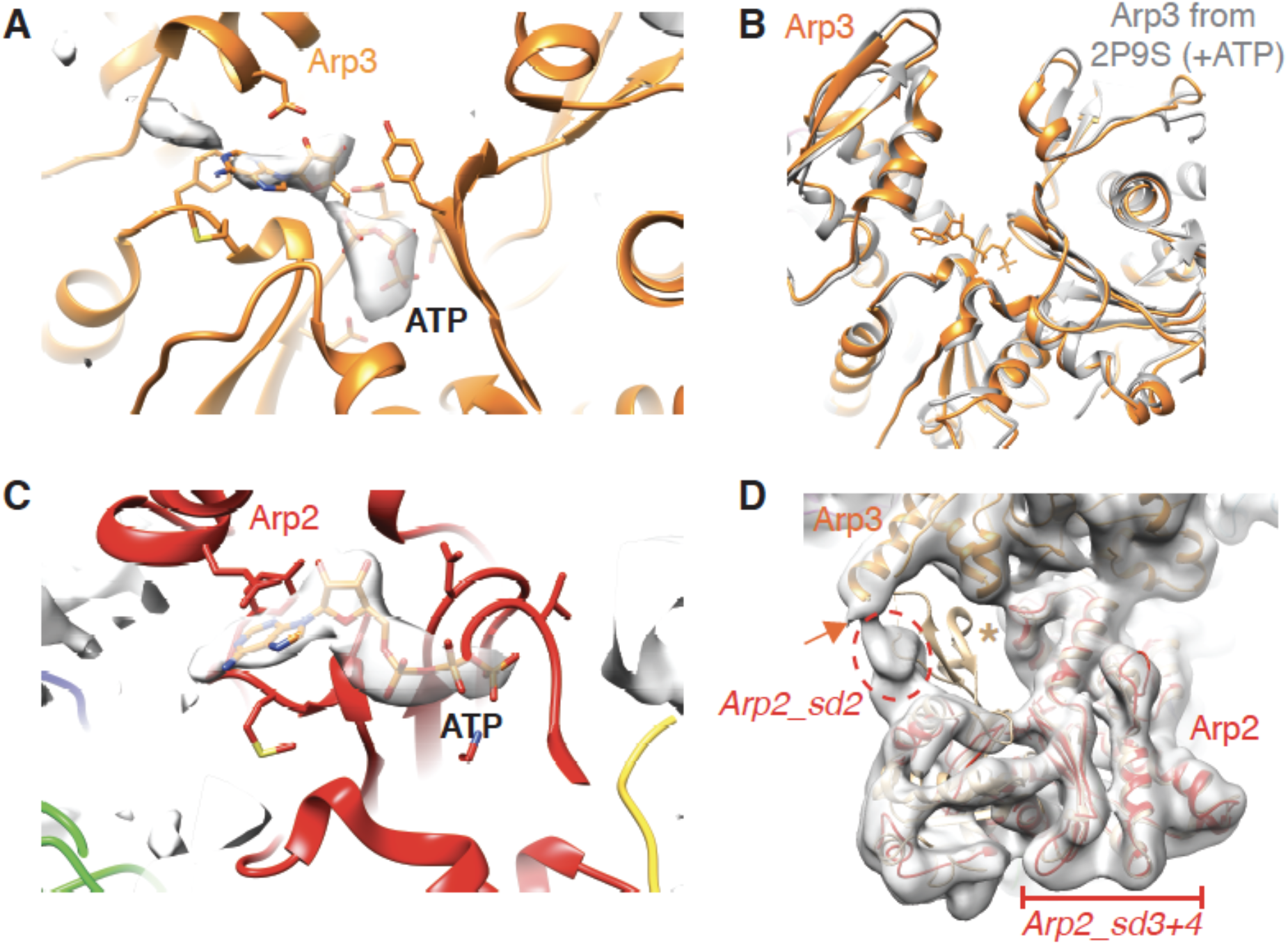
Nucleotide binding sites of Arp3 and Arp2 in Arp2/3-C1B-C5L. (A) Ribbon depiction of the Arp3 model with density corresponding to bound nucleotide shown in surface representation. This density is the calculated difference between our cryo-EM reconstruction and simulated density of the atomic model without nucleotide at equivalent resolution, calculated using Chimera. This supports the conclusion that ATP is bound to Arp3; (B) Conformation of the nucleotide binding pocket of ATP-bound Arp3 aligned (on subdomain 3) with a previously determined structure of ATP-bound Arp2/3 (2P9S; (Nolen and Pollard, 2007)), showing equivalent closure of the pocket in the presence of bound nucleotide compared to the absence of nucleotide (Figure 1D); (C) Ribbon depiction of the Arp2 model with density corresponding to bound nucleotide shown in surface representation. This density is the calculated difference between our cryo-EM reconstruction and simulated density of the atomic model without nucleotide at equivalent resolution, calculated using Chimera. This supports the conclusion that ATP is bound to Arp2. (D) Arp2 in the GMF-inhibited Arp2/3 (PDB 4JD2, in tan) (Luan and Nolen, 2013) is shown aligned with Arp2 (red) in Arp2/3-C1B-C5L within the low-pass filtered cryo-EM density. As previously shown, in the Arp2/3-C1B-C5L reconstruction Arp2 subdomain 2 is flexibly connected to Arp3 helix-α9 (orange arrow), whereas the well-defined structure of the Arp2 subdomain 2 in the GMF-inhibited structure adopts a different conformation which protrudes from the EM density (tan asterisk). For clarity, other subunits within the GMF-inhibited complex are not shown.

**Figure 1 - figure supplement 3.**
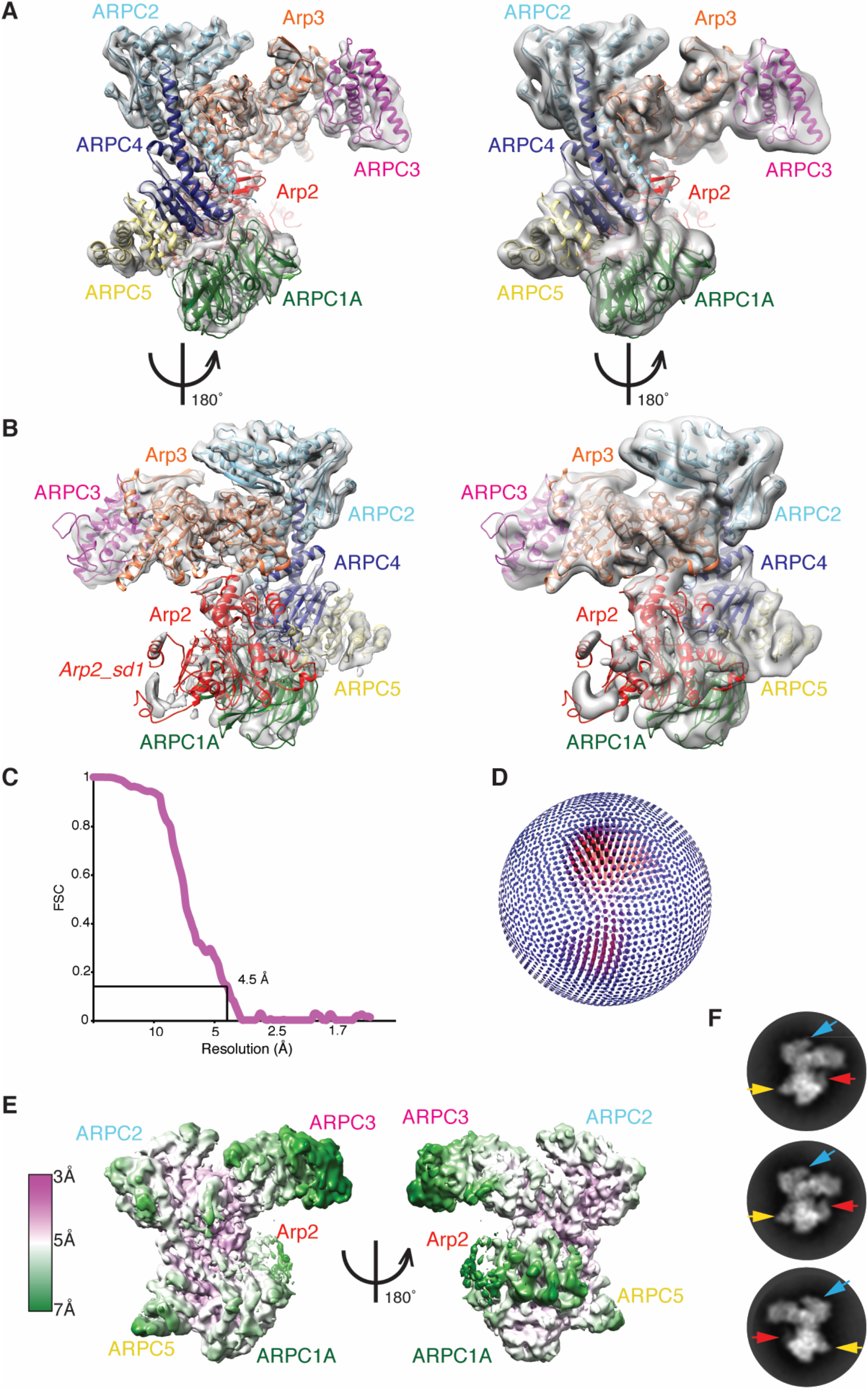
The cryo-EM reconstruction of the human Arp2/3-ARPC1A-ARPC5 complex. (A) Left, cryo-EM reconstruction of Arp2/3-C1A-C5; right, same view with a ~8Å low-pass filter applied to potentially reveal more flexible regions of the complex at lower resolution. The docked model is coloured as in previous figures: Arp2: red; Arp3: orange; ARPC2: cyan; ARPC3: dark pink; ARPC4: blue, except that ARPC1A is dark green and ARPC5 is pale yellow; (B) Left, 180º rotated view compared to (A) of the Arp2/3-C1A-C5 reconstruction and model; right, same view of the reconstruction with a ~8Å low-pass filter applied; (C) FSC curve of the Arp2/3-C1A-C5 reconstruction, showing the 0.143 criteria resolution cut-off = 4.5Å; (D) Orientation distribution of particles used for the final 3D reconstruction; the most common view corresponds to that depicted in panel A; (E) Local resolution depiction of the Arp2/3-C1A-C5 reconstruction calculated using Relion, showing views equivalent to panel A (left) and panel B (right). (F) 2D class averages of Arp2/3-C1A-C5 showing views corresponding to panel A (upper 2 classes) and panel B (bottom panel) illustrating the variable density corresponding to subdomain 2 of Arp2 (red arrows) but the consistent density corresponding to ARPC5 (yellow arrows); this contrasts to that seen in the Arp2/3-C1B-C5L complex. ARPC2 (blue arrow) is also indicated for reference.

**Figure 2 - figure supplement 1.**
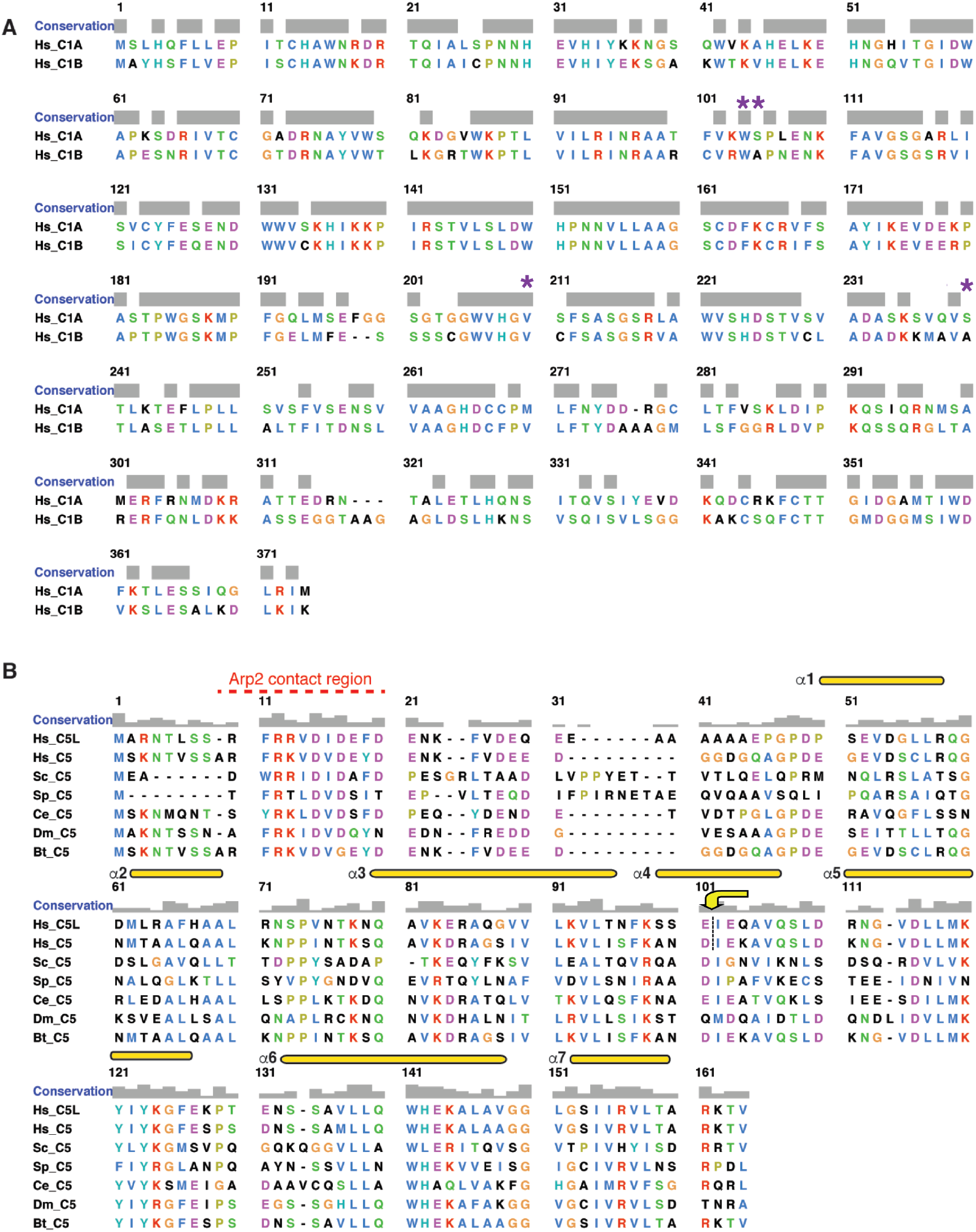
Sequence alignment of ARPC1 and ARPC5. (A) Sequence alignment of human ARPC1 isoforms. Sequences were aligned using T-Coffee (Notredame et al., 2000) and the alignment was prepared using Chimera (Pettersen et al., 2004) with the Clustal X colour scheme. The position of two immunodeficiency syndrome-associated point mutations in ARCP1B – W104S, A105V, V208F and A238T – are indicated with purple asterisks (Kahr et al., 2017). (B) Sequence alignment of ARPC5 including the two human isoforms (Hs_C5L, Hs_C5), and C5 sequences from *S. cerevisiae* (Sc_S5), *S. pombe* (Sp_C5), *C. elegans* (Ce_C5), *D. melanogaster* (Dm_C5) and *B. taurus* (Bt_C5). As, above, sequences were aligned using T-Coffee, the alignment was prepared using Chimera with the Clustal X colour scheme, and the main secondary structural elements are annotated above the alignment according to (Robinson et al., 2001). The yellow arrow/dotted line indicates the splicepoint in the C5/C5L hybrids.

**Figure 2 - figure supplement 2.**
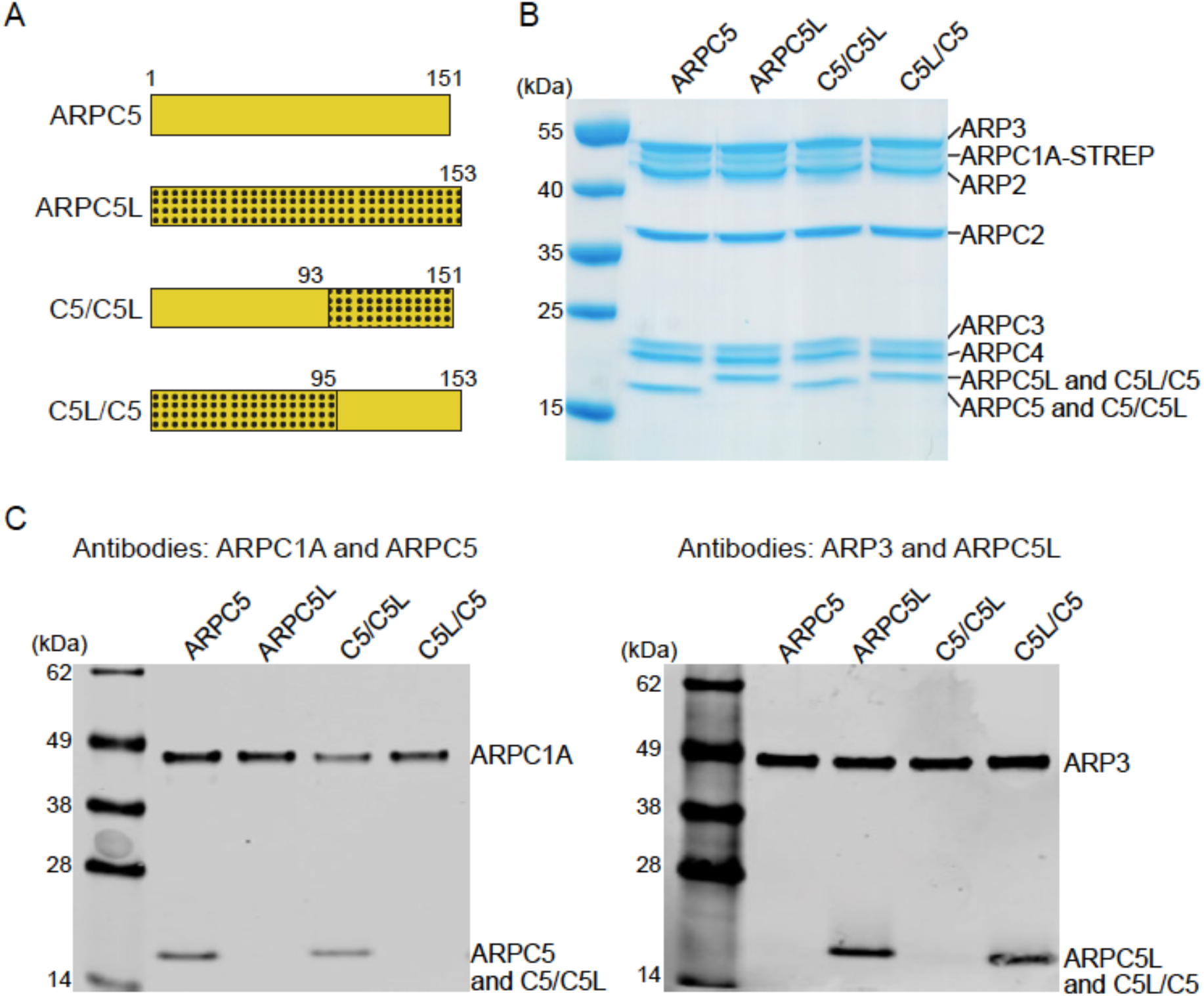
Production and characterization of ARPC5/C5L hybrid complexes. (A) Schematic and nomenclature of ARPC5/C5L hybrids. (B) Coomassie-stained gel of purified recombinant Arp2/3 complexes containing ARPC1A together with ARPC5, ARPC5L or their hybrids. (C) Immunoblot analysis of purified recombinant Arp2/3 complexes used in this study.

